# Intracortical remodelling increases in highly-loaded bone after exercise cessation

**DOI:** 10.1101/2022.05.06.490923

**Authors:** Raniere Gaia Costa da Silva, Tsim Christopher Sun, Ambika Prasad Mishra, Alan Boyde, Michael Doube, Christopher Michael Riggs

## Abstract

Resorption within cortices of long bones removes excess mass and damaged tissue, and increases during periods of reduced mechanical loading. Returning to high-intensity exercise may place bones at risk of failure due to increased porosity caused by bone resorption. We used microradiographs of bone slices from highly-loaded (metacarpal, tibia) and minimally-loaded (rib) bones from 12 racehorses, 6 that died during a period of high-intensity exercise and 6 that had a period of intense exercise followed by at least 35 days of rest prior to death, and measured intracortical canal cross-sectional area (Ca.Ar) and number (N.Ca) to infer remodelling activity across sites and exercise groups. Large canals that are the consequence of bone resorption (Ca.Ar > 0.04 mm^2^) were 1.4× to 18.7× greater in number and area in the third metacarpal bone from rested than exercised animals (p = 0.005– 0.008), but were similar in number and area in ribs from rested and exercised animals (p = 0.575–0.688). An intermediate relationship was present in the tibia, and when large canals and smaller canals that result from partial bony infilling (Ca.Ar > 0.002 mm^2^) were considered together. The mechanostat may override targeted remodelling during periods of high mechanical load by enhancing bone formation, reducing resorption and suppressing turnover. Both systems may work synergistically in rest periods to remove excess and damaged tissue.

## Introduction

Osteoclastic resorption of bone serves several critical functions. It is essential in the mechanism for expression of skeletal adaptation to load: resorption also facilitates the reduction of bone mass as a response to reduced loads on the skeleton, moulds the surfaces of bones to refine their geometric properties in response to altered strain (Andreasen et al., 2020; Bakalova et al., 2018; Ireland et al., 2014; Shaw & Ryan, 2012; Shaw & Stock, 2009) and is the first stage in the process to replace bone with one microstructure (Busse et al., 2010) or fibre orientation (Goldman et al., 2003, 2014; McFarlin et al., 2008; Riggs et al., 1993b, 1993a) with that of another (Collins et al., 2019). It also facilitates the repair of bone tissue by removing damaged or dead bone matrix to make way for its replacement with healthy tissue (Mori & Burr, 1993). Furthermore, resorption of bone is important for mineral homeostasis, providing a mechanism to release mineral ions bound in bone into circulation (Cappariello et al., 2014).

While osteoclastic resorption is an ongoing process in the bones of most mammals some evidence supports focal targeting of resorption in bones of the appendicular skeleton in regions where the matrix has suffered fatigue as a consequence of mechanical overload or repetitive stress injury (Martin, 2007; Plotkin, 2014). Non-targeted or stochastic remodelling may occur as a homeostatic mechanism that has evolved to maintain the material health of the skeleton by reducing the effects of fatigue damage on bones in the long term (Parfitt, 2002). The suppression of both targeted and stochastic remodelling by experimental administration of bisphosphonates provides support for this hypothesis (Li et al., 2001). Resorption, the essential first stage of the remodelling process in dense bone, creates additional porosity that may relate to higher strain energy density (Bakalova et al., 2018), decreased elastic modulus (Gibson et al., 2006) and diminished toughness (Bell et al., 1999; Schaffler & Burr, 1988; Yeni et al., 1997).

The loss of strength and stiffness caused by increased porosity may predispose bone to accelerated fatigue or monotonic failure in the face of ongoing cyclical loads (Frank et al., 2002; Seref-Ferlengez et al., 2015). Bone must strike a balance between the risks of tissue deterioration occurring through the accumulation of damage with the risks of failure due to increased porosity incurred during remodelling. Previous studies indicate that the balance falls in favour of allowing damage to accumulate (Seref-Ferlengez et al., 2015) rather than attempting repair by remodelling while high loads continue.

Horses are a unique model species in which to study bone dynamics. Their bones are large and support plentiful secondary osteonal remodelling (Boyde, 2003; Firth et al., 2005; Riggs et al., 1993a; Shahar et al., 2011; Stover et al., 1992), which does not normally occur in laboratory rodents or other mammals smaller than about 2 kg body mass (Felder et al., 2017). Material for detailed histological study is available post-mortem from equine athletes with greater regularity and control than from human athletes. Histological examination of bone from horses used in exercise tests on treadmills (Firth et al., 2005) and from post mortem specimens from racehorses (Whitton et al., 2013) suggests that osteoclastic resorption in the third metacarpal bone is suppressed in horses that are consistently subjected to a high exercise workload compared to those forced to rest. A physiological control system may have evolved to favour the safest way to manage skeletal integrity in the face of ongoing mechanical loading, but evidence for this hypothesis is weak. The purpose of the present study was to develop the evidence base to better understand bone’s response to periods of intense exertion and subsequent rest.

In this study we measured bone remodelling, using the number and size of resorption canals as a proxy, in selected regions of bones from horses in active training and from horses subject to enforced rest. We used samples taken from horses for which we knew the exercise history with high confidence because all the animals resided, trained and raced within one organisation that kept records of all interventions.

## Materials and Methods

Bone was obtained post-mortem from Thoroughbred racehorses (*n*=12) at the Hong Kong Jockey Club (HKJC) that had been euthanised for reasons unrelated to the study (Table 1). Permission to use residual tissues is granted by prior agreement with owners of all horses at the Hong Kong Jockey Club. Racing and training information was taken from the HKJC Racing information Database (The Hong Kong Jockey Club, 2021). Horses were assigned to one of two groups: “exercised” (*n*=6) and “rested” (*n*=6). The “exercised” group was restricted to horses that had been in regular training for racing and had undertaken intense exercise within seven days of their death. “Intense exercise” was defined as a race, barrier trial (practice race), training gallop or training canter and “regular” as this level of exercise occurred on at least four occasions in the preceding 30 days. The “rested” group was restricted to horses that had previously been in race training (at which stage they had undertaken similar intense exercise to that of the “exercised” group) but that were undergoing a period of rest and had not undertaken any intense exercise for 30–120 days prior to euthanasia.

**Table 1.** List of horses in the study.

| Group | ID | Age (years) | Intense exercise (days) | Exercise-death interval (days) | Cause of death |
| --- | --- | --- | --- | --- | --- |
| Exercised | 01 | 5 | 8 | 0 | Drug anaphylaxis |
|  | 02 | 6 | 8 | 0 | Tendon rupture |
|  | 03 | 6 | 6 | 3 | Collapse during racing |
|  | 04 | 5 | 4 | 1 | colic |
|  | 05 | 5 | 6 | 0 | Collapse during racing |
|  | 06 | 4 | 6 | 0 | Collapse during fast work |
|  | mean | 5.17 | 6.33 | 0.67 |  |
|  | Std. Dev. | 0.75 | 1.51 | 1.21 |  |
| Rested | 07 | 8 | 5 | 35 | Advanced osteoarthritis |
|  | 08 | 6 | 5 | 91 | Tendon rupture |
|  | 09 | 8 | 5 | 82 | Chronic lameness |
|  | 10 | 8 | 4 | 45 | Advanced osteoarthritis |
|  | 11 | 5 | 5 | 79 | Unsuitable temperament |
|  | 12 | 6 | 6 | 94 | Chronic lameness |
|  | mean | 6.83 | 5.00 | 71.0 |  |
|  | Std. Dev. | 1.33 | 0.63 | 24.8 |  |

Bone sections were cut from the right third metacarpal bone and right tibia, at locations that are common sites for fatigue fractures in Thoroughbred racehorses (Riggs & Pilsworth, 2014). Sections were also cut from the mid-body of the tenth left rib to represent bone of the axial skeleton that is subject to loads that increase significantly less than loads on the limbs during locomotion (Figure 1).

**Figure 1.**
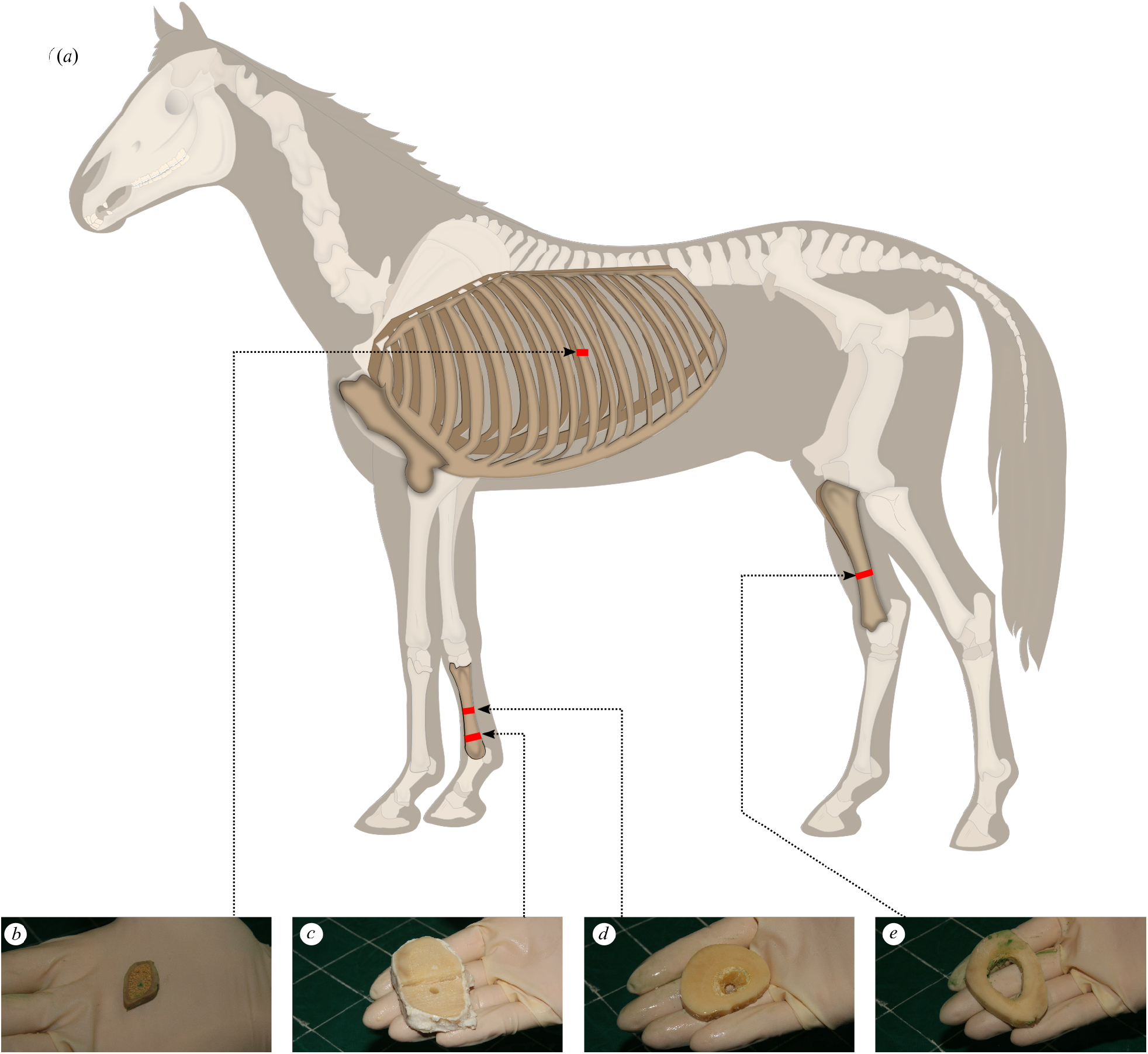
Sample collection sites mapped to (a) the whole horse: (b) mid-diaphysis of the left tenth rib (c) lateral distal metaphysis of the right third metacarpal bone; (d) mid-diaphysis of the right third metacarpal bone; (e) distal third region of the right tibia. This work is a derivative of “Horse anatomy.svg” by Wikipedian Prolific and Wilfredor used under Creative Commons Attribution-Share Alike 3.0 Unported licence.

The bones were dissected in their entirety immediately post mortem. Transverse blocks of 20 mm thickness were cut from the specified region of each bone using a band saw (ST-WBS 180). The blocks from the rib, tibia and mid diaphysis of third metacarpal included the entire cross section of the bone. The blocks of the distal metaphysis of the third metacarpal included only the lateral half of the entire cross section. Blocks were dissected free of soft tissue including fat and periosteum before being placed in a bacterial pronase detergent (Tergazyme 10% alkaline pronase detergent, Alconox Inc, NY, USA) at 37 °C for five days to digest remaining soft tissue. Blocks were then fixed in 70% ethanol for seven days. Plane-parallel sections 250 μm and 100 μm thick were subsequently cut from each block in a plane at right angles to the long axis of the bone using an annular saw (Leica Microsystems SP1600). Sections were stored in glycerol (Figure 2). The blocks of the distal third region of the right tibia and of the mid-diaphysis of the right third metacarpal bone of horse 11 were severely damaged during cutting and could not be used.

**Figure 2.**
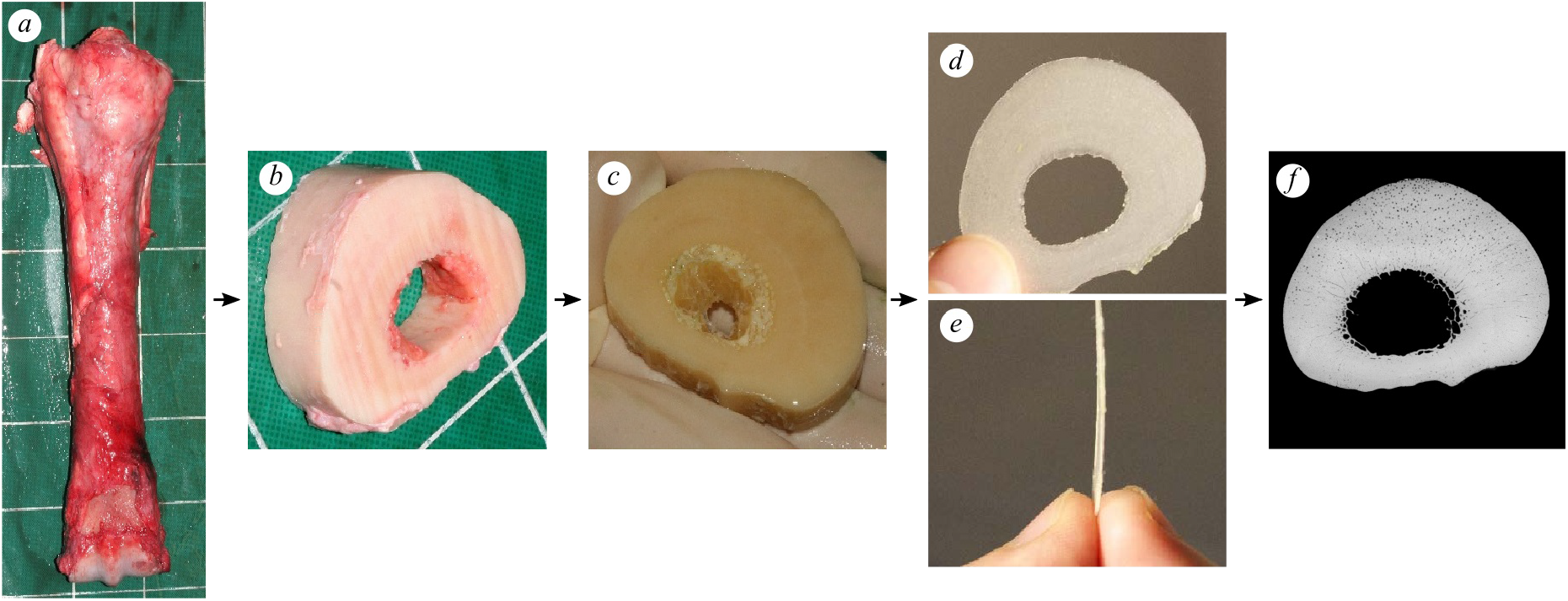
Bone processing from post-mortem collection to microradiograph: (a) dorsal view of the right third metacarpal bone immediately post mortem, (b) transverse 20 mm thick slice cut with a band-saw, (c) slice after five days of immersion in bacterial pronase detergent and subsequent fixation for seven days in 70\% ethanol, (d) proximodistal and (e) lateral view of 250 μm thick sections cut with an annular saw, (f) microradiograph of 250 μm thick section. Lateral to the left, dorsal to the top (c, d, f).

Microradiographs (Figure 3) were obtained in 16-bit DICOM format from each section by point projection digital microradiography (Faxitron, QADOS, Sandhurst, Berkshire, UK) at 26 kV and the maximum magnification possible so that each section was captured in its entirety by a single image (Table 2, Figure 3). Images were converted to 8-bit TIFFs for subsequent analysis and these are available from figshare (see Supplementary material).

**Table 2.** X-ray magnification, pixel spacing and minimum resolvable canal diameter (Ca.D) information for each sample site. Magnification was the maximum that allowed the entire specimen to be imaged in a single frame.

| bone section | magnification | pixel spacing<br>($\mu\text{m}$) | minimum Ca.D<br>($\mu\text{m}$) |
| --- | --- | --- | --- |
| rib | 5 | 10.0 | 11.28 |
| tibia | 2 | 25.0 | 28.21 |
| metacarpal diaphysis | 2 | 25.0 | 28.21 |
| metacarpal metaphysis | 3 | 16.7 | 18.84 |

**Figure 3.**
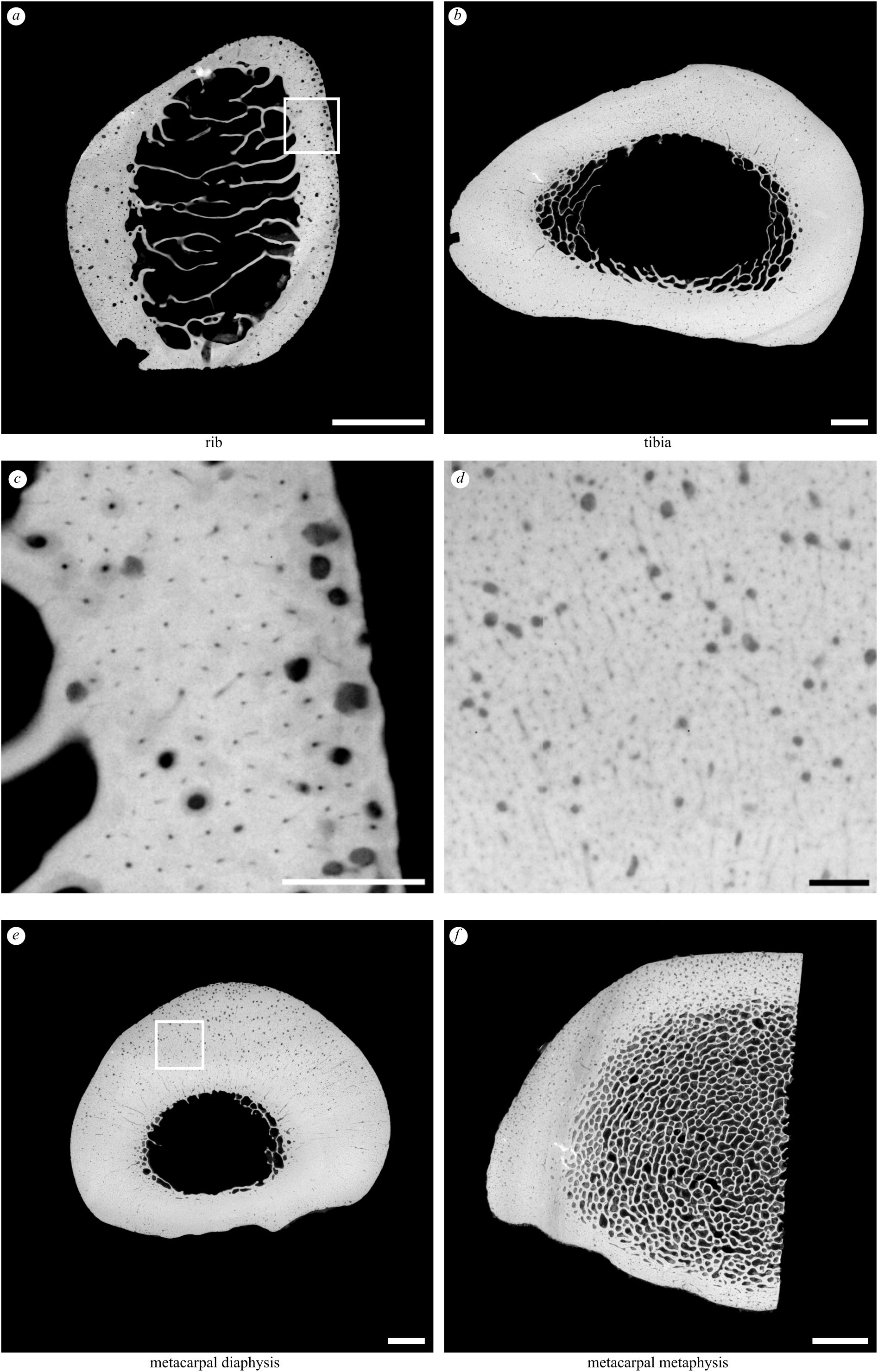
Example microradiographs obtained from each section: (a) transverse section of the mid-diaphysis of the left tenth rib (5× magnification); (b) transverse section of the distal third region of the right tibia (2× magnification); (e) transverse section of the mid-diaphysis of the right third metacarpal (2× magnification); (f) transverse section of the lateral half of the distal metaphysis of the right third metacarpal (3× magnification). Box in (a) magnified in (c); box in (e) magnified in (d). Scale bars: a, b, e, f, 5 mm; c, d, 1 mm. Lateral to left, cranial/dorsal to top.

Microradiographs of each section were analysed using Fiji 1.53c (Rueden et al., 2017; Schindelin et al., 2012; Schneider et al., 2012) to quantify nonmineralised areas of the section (porosities) that appear as small radiolucent spots surrounded by radiopaque cortical bone matrix. Cortical bone was segmented by manually drawing along the endosteal and periosteal boundaries. A triangle method threshold (Zack et al., 1977) was calculated for each image and applied to the greyscale microradiographs, to assign each pixel to one of ‘bone’ or ‘pore’ binary phases. ImageJ’s Analyze Particles function was used to identify individual pores (Figure 4c,d,e). Particles were filtered using Python to retain those with circularity between 0.3 and 1 and area > 0.002 mm^2^. Size categories were selected to coincide with diameters associated with mature Haversian canals (Ca.Dm, 20 – 30 μm in the horse) and varying stages of resorption/infilling of resorption canals (Felder et al., 2017). The smallest particles with canal area (Ca.Ar) ≤ 0.002 mm^2^ were assumed to represent mature Haversian canals and were excluded from further analysis. Large, smooth pores at the endosteal margin of the cortex in each section that reflected transition to cancellous bone were manually excluded. Remaining pores were assumed to represent regions of bone resorption with or without subsequent formation, with the most likely descriptions corresponding to osteonal canals at the cutting cone (Ca.Ar > 0.04 mm^2^) and closing cone (Ca.Ar = 0.002-0.04 mm^2^) stages (Figure 4). The diameter of each canal (Ca.Dm, μm) was calculated from Ca.Ar assuming a circular cross section and recorded. The cutoff values of 0.002 and 0.04 mm^2^ were calculated from canal diameters of 50 and 225 μm (0.05 and 0.225 mm) as 0.002 mm^2^ = *π* × (0.025 mm)^2^ and 0.04 mm^2^ = *π* × (0.1125 mm)^2^ using A = *πr*^2^, where *r* is the radius (half the diameter) of the canal. The minimum canal diameter (50 μm) was twice that of the largest pixel spacing in the images (Table 2). Cortical bone area (Ct.Ar, μm^2^), count of individual canals (N.Ca, unitless), total canal area (Tt.Ca.Ar, μm^2^), canal numerical density (N.Ca/Ct.Ar, μm^−2^), and canal area fraction (Tt.Ca.Ar/Ct.Ar, unitless), were calculated for each section (Dempster et al., 2013). Differences in the measurements between exercised and rested groups were tested for using the Mann-Whitney U test, implemented in SciPy (SciPy 1.0 Contributors et al., 2020). Full details of the image processing and statistical analysis are available at figshare (See Supplementary material).

**Figure 4.**
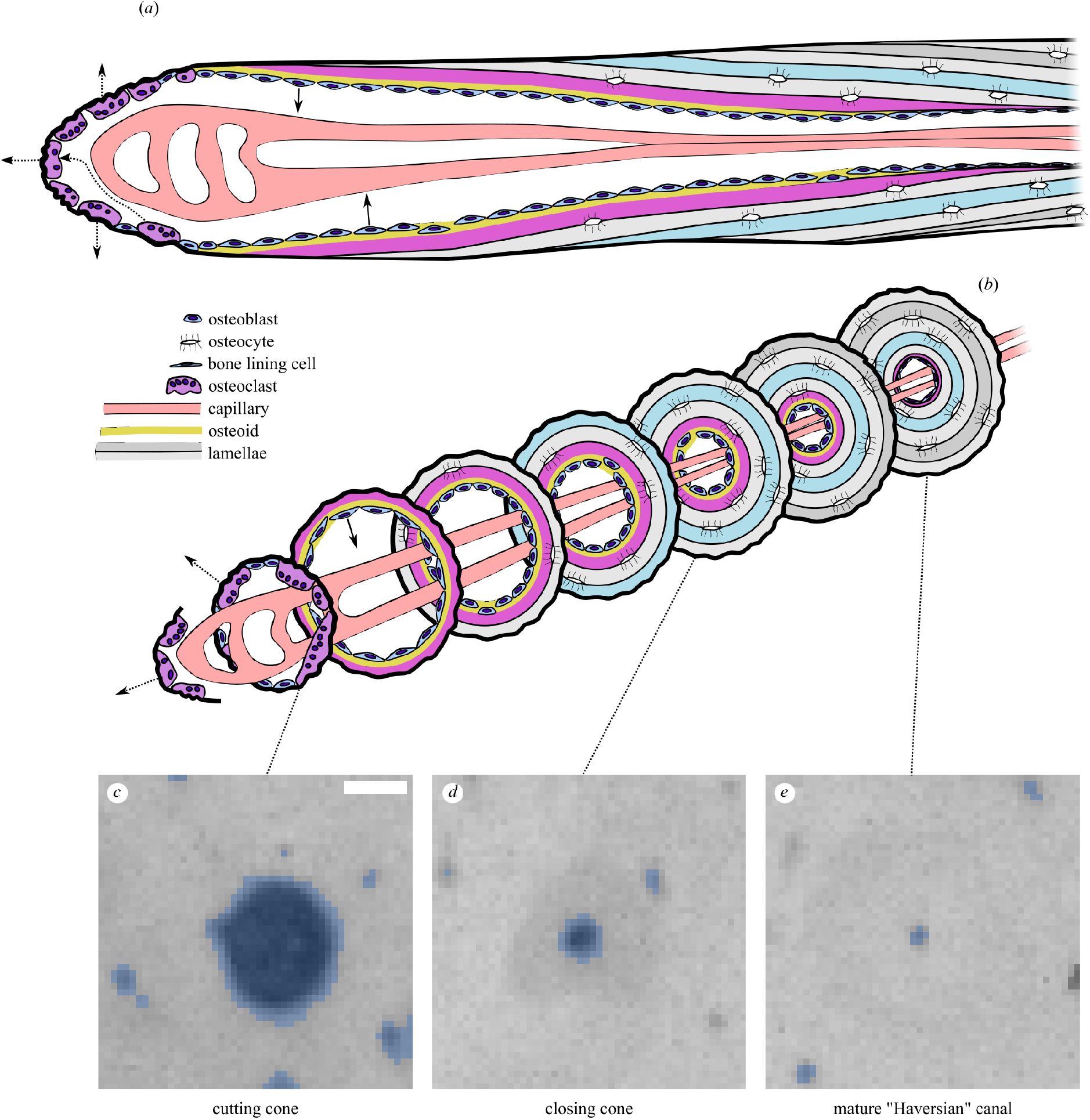
Illustration of secondary osteonal remodelling “basic multicellular unit” in cortical bone in (a) longitudinal section and (b) a series of transverse sections along its length. Representative microradiographs of the (c) cutting cone, (d) closing cone and (e) mature Haversian canal, shown with binary masks derived from processing with Fiji’s Analyze Particles overlaid in blue. In this study we classify canals into large (Ca.Ar > 0.04 mm^2^; (c)) and combined large and small (Ca.Ar > 0.002 mm^2^; (c+d)), and ignore the smallest canals (Ca.Ar ≤ 0.002 mm^2^; (e)). Scale bar 100 μm. Modified from (Doube, 2022) under CC-BY licence terms.

## Results

Where groups differed, samples from the rested group contained more canals than samples from the exercised group (Figure 5, 6; Tables 3, 4, 5, 6), which was most pronounced in the metacarpal sites and for the largest canals. Numbers of canals (N.Ca/Ct.Ar) and overall porosity (Tt.Ca.Ar/Ct.Ar) were 1.4× – 18.7× greater in rested than exercised horses’ metacarpal diaphysis and metacarpal metaphysis, but were not different between rested and exercised horses in the rib or tibia (Table 3, 5). Large canals (Ct.Ar > 0.04 mm^2^) were unevenly distributed in some specimens, appearing more densely in the dorsal cortex of the rested metacarpal diaphysis (Figure 5g) and the caudal tibia (Fig. 5f), while a higher number of diffusely-located large canals was evident in the rested metacarpal metaphysis (Fig. 5h).

**Table 3.** Summary of canal dimensions by site and exercise group (mean ± s.d.) for cutting and closing cones (Ca.Ar > 0.002 mm^2^), normalised by cortical area (Ct.Ar; identical to Table 5 because they are the same specimens).

| bone | group | Ct.Ar<br>($\text{mm}^2$) | N.Ca | Tt.Ca.Ar<br>($\text{mm}^2$) | N.Ca/Ct.Ar<br>($1/\text{mm}^2$) | Tt.Ca.Ar/Ct.Ar |
| --- | --- | --- | --- | --- | --- | --- |
| rib | exercised | $80.8 \pm 5.0$ | $232 \pm 102$ | $1.36 \pm 0.97$ | $2.82 \pm 1.09$ | $0.016 \pm 0.011$ |
| | rested | $79.6 \pm 6.6$ | $260 \pm 130$ | $1.49 \pm 0.80$ | $3.27 \pm 1.65$ | $0.019 \pm 0.010$ |
| tibia | exercised | $824 \pm 20$ | $1527 \pm 678$ | $9.26 \pm 4.79$ | $1.86 \pm 0.82$ | $0.011 \pm 0.006$ |
| | rested | $898 \pm 81$ | $1976 \pm 574$ | $17.4 \pm 4.12$ | $2.21 \pm 0.67$ | $0.019 \pm 0.004$ |
| metacarpal diaphysis | exercised | $783 \pm 73$ | $1392 \pm 561$ | $9.06 \pm 3.33$ | $1.77 \pm 0.69$ | $0.012 \pm 0.005$ |
| | rested | $816 \pm 59$ | $1552 \pm 850$ | $17.4 \pm 5.80$ | $1.92 \pm 1.11$ | $0.021 \pm 0.007$ |
| metacarpal metaphysis | exercised | $254 \pm 27$ | $505 \pm 84$ | $3.03 \pm 0.86$ | $2.03 \pm 0.49$ | $0.012 \pm 0.004$ |
| | rested | $250 \pm 37$ | $730 \pm 171$ | $11.4 \pm 3.52$ | $2.91 \pm 0.58$ | $0.046 \pm 0.011$ |

**Table 4.** Results of Mann-Whitney comparisons between exercised and rested groups for canals with Ca.Ar > 0.002 mm^2^. *U* values tend towards 0 as the overlap between the two groups’ rank distributions decreases to 0. All groups have *n* = 6 except rested tibia and rested metacarpal diaphysis which have *n* = 5.

| bone | Ct.Ar |  | N.Ca |  | Tt.Ca.Ar |  | N.Ca/Ct.Ar |  | Tt.Ca.Ar/Ct.Ar |  |
| --- | --- | --- | --- | --- | --- | --- | --- | --- | --- | --- |
| | $U$ | $p$ | $U$ | $p$ | $U$ | $p$ | $U$ | $p$ | $U$ | $p$ |
| rib | 21 | 0.69 | 15 | 0.69 | 16 | 0.81 | 17 | 0.94 | 14 | 0.58 |
| tibia | 6 | 0.12 | 8 | 0.24 | 4 | 0.06 | 10 | 0.41 | 5 | 0.08 |
| metacarpal diaphysis | 12 | 0.65 | 16 | 0.93 | 3 | 0.04 | 16 | 0.93 | 4 | 0.06 |
| metacarpal metaphysis | 19 | 0.94 | 6 | 0.07 | 0 | 0.01 | 5 | 0.05 | 0 | 0.01 |

**Table 5.** Summary of canal dimensions by site and exercise group (mean ± s.d.) for cutting cones (Ca.Ar > 0.04 mm^2^), normalised by cortical area (Ct.Ar; identical to Table 3 because they are the same specimens).

| bone | group | Ct.Ar (mm<sup>2</sup>) | N.Ca | Tt.Ca.Ar (mm<sup>2</sup>) | N.Ca/Ct.Ar (1/mm<sup>2</sup>) | Tt.Ca.Ar/Ct.Ar |
| --- | --- | --- | --- | --- | --- | --- |
| rib | exercised | 80.8 ± 5.0 | 3.67 ± 4.41 | 0.20 ± 0.25 | 0.04 ± 0.05 | 0.002 ± 0.003 |
| | rested | 79.6 ± 6.6 | 5.00 ± 6.63 | 0.26 ± 0.33 | 0.06 ± 0.08 | 0.003 ± 0.004 |
| tibia | exercised | 824 ± 20 | 165.8± 15.8 | 0.92 ± 0.98 | 0.02 ± 0.02 | 0.001 ± 0.001 |
| | rested | 898 ± 81 | 50.2 ± 26.0 | 2.98 ± 1.87 | 0.06 ± 0.03 | 0.003 ± 0.002 |
| metacarpal diaphysis | exercised | 783 ± 73 | 17.0 ± 9.25 | 1.06 ± 0.63 | 0.02 ± 0.01 | 0.0014 ± 0.0009 |
| | rested | 816 ± 59 | 93.0 ± 44.1 | 5.47 ± 2.49 | 0.11 ± 0.05 | 0.007 ± 0.003 |
| metacarpal metaphysis | exercised | 254 ± 27 | 3.67 ± 3.50 | 0.24 ± 0.20 | 0.014 ± 0.014 | 0.0009 ± 0.0008 |
| | rested | 250 ± 37 | 66.0 ± 41.6 | 3.89 ± 2.59 | 0.26 ± 0.14 | 0.015 ± 0.009 |

**Table 6.** Results of Mann-Whitney comparisons between exercised groups for canals with Ca.Ar > 0.04 mm^2^. *U* values tend towards 0 as the overlap between the two groups’ rank distributions decreases to 0. All groups have *n* = 6 except rested tibia and rested metacarpal diaphysis which have *n* = 5.

| bone | Ct.Ar |  | N.Ca |  | Tt.Ca.Ar |  | N.Ca/Ct.Ar |  | Tt.Ca.Ar/Ct.Ar |  |
| --- | --- | --- | --- | --- | --- | --- | --- | --- | --- | --- |
|  | *U* | *p* | *U* | *p* | *U* | *p* | *U* | *p* | *U* | *p* |
| rib | 21 | 0.689 | 16 | 0.806 | 15 | 0.688 | 14 | 0.575 | 15 | 0.688 |
| tibia | 6 | 0.121 | 4 | 0.055 | 5 | 0.083 | 4 | 0.055 | 5 | 0.083 |
| metacarpal diaphysis | 12 | 0.648 | 0 | 0.008 | 0 | 0.008 | 0 | 0.008 | 0 | 0.008 |
| metacarpal metaphysis | 19 | 0.936 | 0 | 0.005 | 0 | 0.005 | 0 | 0.005 | 0 | 0.005 |

**Figure 5.**
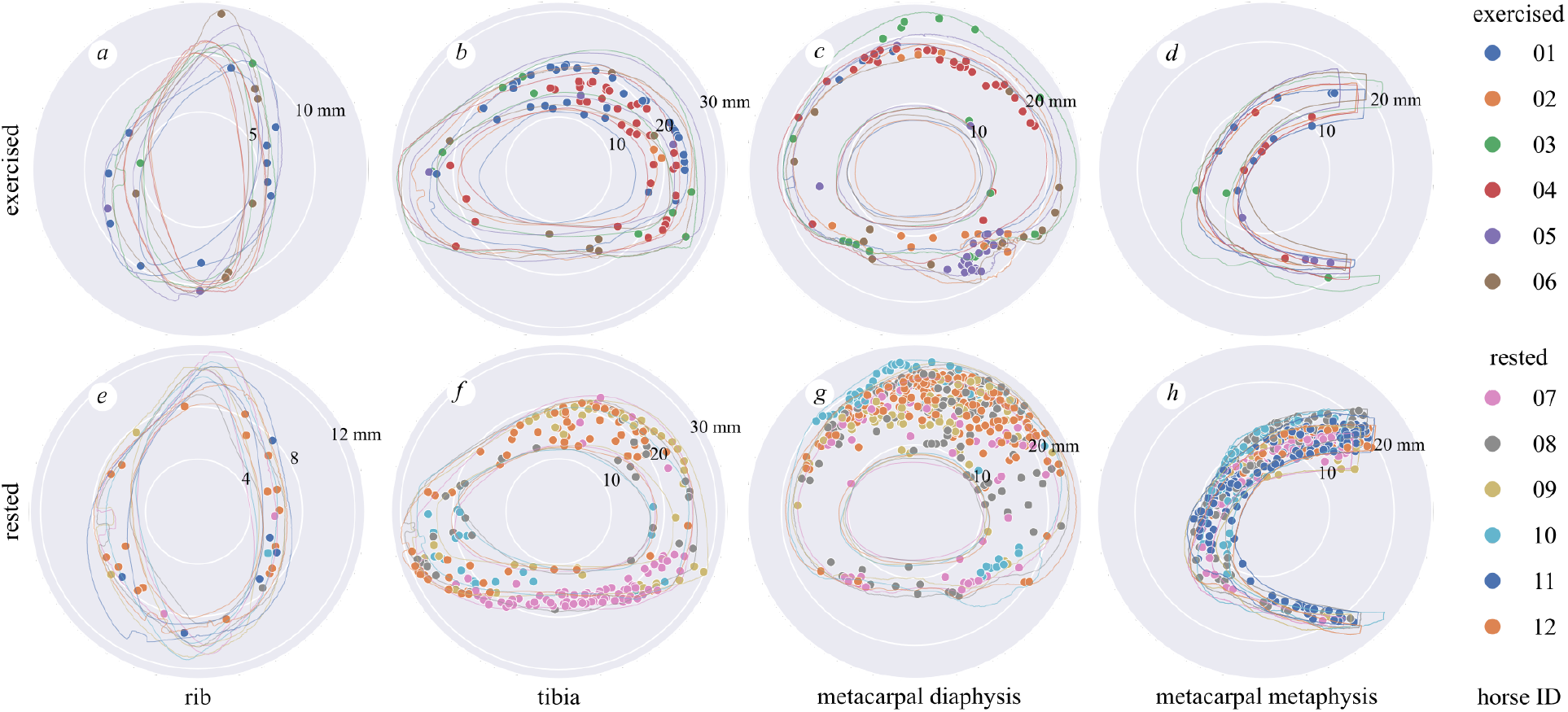
Polar plots of the largest canals representing resorption spaces or cutting cones (Ca.Ar > 0.04 mm^2^) in bones from the exercised (a, b, c, d) and rested (e, f, g, h) groups, centred on each sample’s centre of mass. Canals are indicated by solid dots, and periosteal and endosteal contours are indicated by outlines, all colour coded by horse ID. Note the larger number of large canals in the rested metacarpal and tibial sites and their anatomical distribution in the caudal tibia, dorsal metacarpal diaphysis and throughout the metacarpal metaphysis. Cranial / dorsal to the top, medial to the right. Scale in mm given by numbers indicating the radius of the white circles on each plot. See Tables 5, 6 for statistical summaries and comparisons.

**Figure 6.**
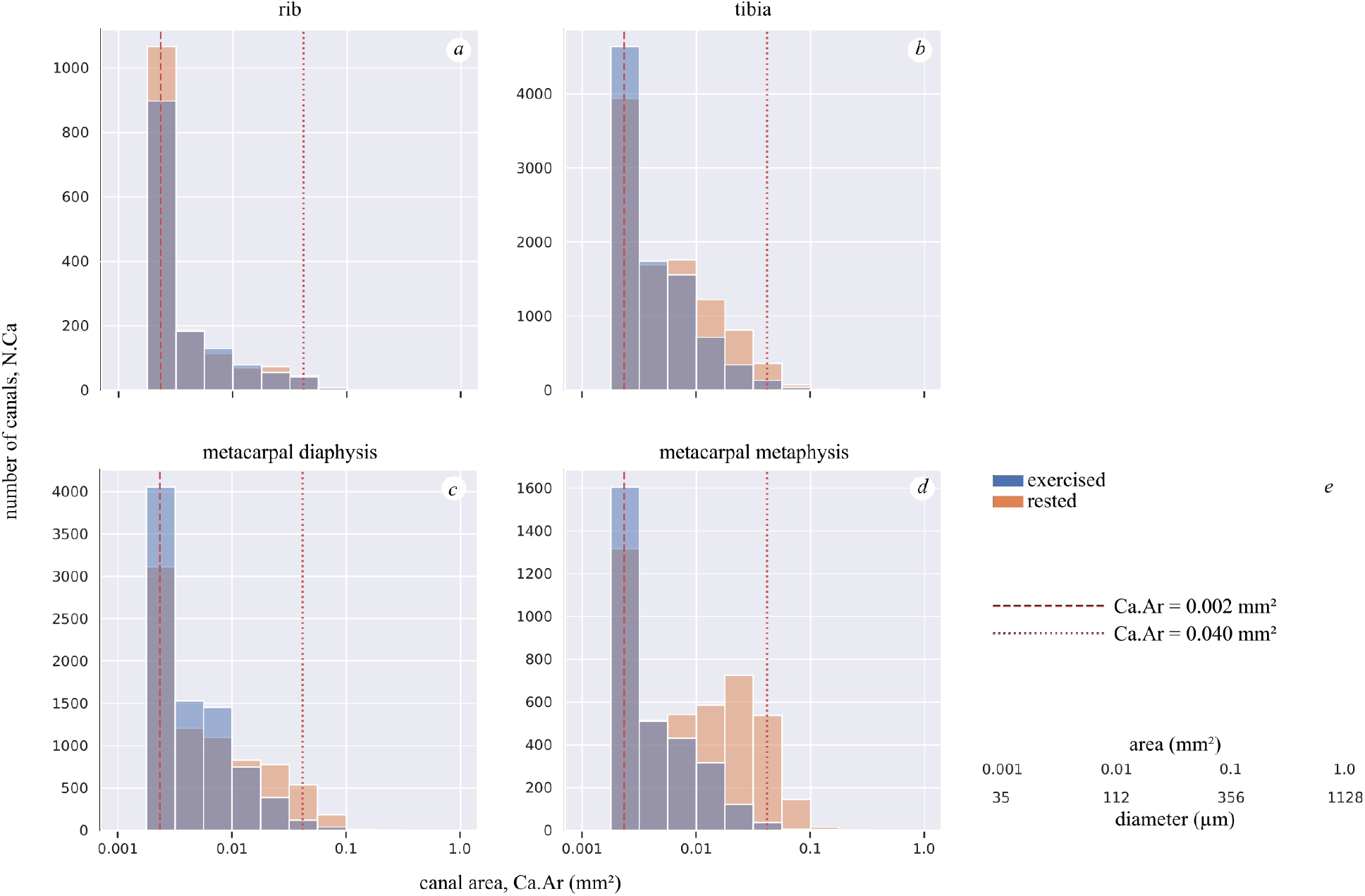
Canal number (N.Ca) summed over all samples within exercise and rested groups versus canal area (Ca.Ar) for canals with Ca.Ar > 0.002 mm^2^, for each sample site. Note the greater number of large canals in the rested metacarpal metaphysis (d), and the similar distributions of canal sizes in rested and exercised ribs (a). See Tables 3, 4, 5, 6 for statistical summaries and comparisons. Vertical dashed line at 0.04 mm^2^ indicates the cutoff value between large (resorption spaces; illustrated in Figure 5) and small (partially infilled) canals. Vertical dashed line at 0.002 mm^2^ indicates the cutoff value between completely infilled mature Haversian canals and closing cones. Conversion between area and diameter assuming a circle provided in (e).

## Discussion

We found significantly more and larger canals of a diameter that matched secondary osteonal remodelling events (i.e. the wide bore of cutting and closing cones) in the cortices of limb bones in resting horses than in horses undergoing high-intensity exercise. Assuming that the canals reflect resorptive activity, the finding provides further evidence to support the hypothesis that a physiological mechanism has evolved to limit or suppress resorption in the face of ongoing, high cyclical loads on bone (Dittmer & Firth, 2017; Komori, 2013), even while damage is accumulating. Because resorption is the first stage of intracortical remodelling, targeted remodelling of fatigue damaged tissue may be reduced, and not increased as suggested by others (Hughes et al., 2020), while intense exercise is ongoing. Here, the rest period may have removed the suppression of resorption, allowing targeted remodelling to proceed. No relationship between osteonal remodelling and exercise group was noted in the rib, which is likely to experience only mildly elevated mechanical loads associated with increased respiratory effort at exercise.

An important function of osteoclastic resorption of bone is the repair of bone, through the replacement of damaged matrix with new, healthy tissue. Evidence that osteoclasts are recruited preferentially to target areas of matrix damage or osteocyte necrosis in cortical bone supports this concept (Martin, 2007; Parfitt, 2002; Plotkin, 2014). Accumulation of fatigue damage probably has little impact on mechanical properties of cortical bone until it becomes severe (Norman et al., 1998; Parfitt, 2002; Plotkin, 2014; Seref-Ferlengez et al., 2015), while suppression of remodelling by half may result in three times the accumulation of fatigue damage (Allen et al., 2006; Mashiba et al., 2000). In some cases, microdamage may improve the toughness of bone (Nalla et al., 2005; Ural & Vashishth, 2007). Bone repair without resorption may occur by organic and inorganic substrates bridging and sealing cracks (Boyde, 2003; Seref-Ferlengez et al., 2014), which might be an inherent property of mineralised tissues occurring across all vertebrate taxa (Herbst et al., 2019). In vitro mechanical testing of bone specimens machined from the third metacarpal of Thoroughbred racehorses showed that the elastic modulus and yield strength of specimens were not significantly affected by prior fatigue loading equivalent to a lifetime of racing (Martin et al., 1996). Conversely, there is good evidence that cortical porosity is negatively correlated with Young’s modulus (Schaffler & Burr, 1988), compressive ultimate stress (Behrens et al., 1974), and fracture toughness (Ural & Vashishth, 2007). Changes in porosity account for more than 75% of the variability in the strength of cortical bone (McCalden et al., 1993), and fatigue-induced damage located near cortical pores may be more likely to lead to fracture than that located in regions of high mineral content (Turnbull et al., 2014).

In the present study, resorption was determined by mapping porosities with a diameter greater than 50 μm (area greater than 0.002 mm^2^). Porosities in this size range were assumed to represent resorption spaces and partially-infilled canals, whose number and size may be an accurate histological measurement of bone remodelling (Frost, 1969; Parfitt, 2002). Porosities within cortical bone are typically present in the form of osteocyte lacunae (<5 μm diameter) with associated, tortuous interconnected canaliculi (<1 μm diameter); Haversian canals at the centre of osteons (c. 20 μm diameter); Volkmann’s canals, running roughly perpendicular to Haversian canals; resorption canals, in varying stages of being infilled with new bone (diameter between 20 and 400 μm). Resorption spaces in cortical bone may be the classical cutting cone passing through bone matrix, or resorption occurring on the surface of existing osteonal canals (Andreasen et al., 2018, 2020). Haversian and resorption canals are roughly circular to ellipsoidal in cross section. The former two predominantly lie parallel with the long axis of the bone and so appear circular in transverse section of long bones. Conversely Volkmann’s canals appear as tubular or elongated oval structures, depending on their precise orientation relative to the plane of section. Larger (>600 μm wide), irregular porosities are present close to the endosteal surface of long bones, representing a transition to trabecular bone (Andreasen et al., 2020): we manually excluded these during segmentation.

Subperiosteal primary osteonal bone growth may occur and osteonal canals rapidly infill early in training in a site-specific manner (Firth et al., 2005), conforming to the mechanostat model whereby increased strains lead to net bone deposition (Frost, 2003). Our data show decreased resorption (fewer large canals) during exercise, and increased resorption (more large canals) during rest following exercise, in bone regions such as the dorsal metacarpal diaphysis that are known to experience high strains and strain rates that increase with increasing locomotor speed (Davies, 2005, 2006). Some resorption, indicated by a small number of large canals, may occur even in the most highly-loaded regions during exercise periods (Figure 5), consistent with previous findings (Whitton et al., 2013).

Complete fracture of bones of the appendicular skeleton as a consequence of extension of stress fractures is relatively common in racehorses and represents a serious threat to horse welfare (Carrier et al., 1998; Riggs et al., 1999; Samol et al., 2021; Vallance et al., 2013). Better understanding of the natural biology of bone, especially in relation to repair of microdamage and how the mechanical integrity of the bone may be undermined by the repair process, is necessary in order to develop management strategies to mitigate risk. The high number of large canals observed in the rested group has important implications for physical activity management (Armstrong et al., 2004; Hughes et al., 2014; Jacobs et al., 2014; Wik et al., 2020), in particular the re-introduction of training after returning from a prolonged rest period. High porosity may be related to reduced bone yield stress and stiffness (McCalden et al., 1997; Schaffler & Burr, 1988; Wachter et al., 2002) and may also create stress risers allowing fracture propagation (Hernandez & Keaveny, 2006). Physical readiness training, reduced marching, exercise on flat rather than hilly terrain, and adhering to an exercise protocol, reduces stress fractures of the lower limb bones in human military recruits (Chalupa et al., 2016; Milgrom & Finestone, 2017).

There is epidemiological evidence that racehorses returning after an eight week rest period are at greater risk of sustaining complete fractures of the humerus, scapula and tibia (Carrier et al., 1998; Samol et al., 2021; Vallance et al., 2013). It remains to be seen whether a gradual programme of exercise reintroduction is more bone-safe. The time taken for completion of bone remodelling is unknown, however, osteoclasts proceed longitudinally through bone at a rate of about 40 μm d^−1^ (Jaworski & Lok, 1972), while osteoblastic infilling starts at the same time radial resorption ends (Lassen et al., 2017) and at 1.0–1.5 μm d^−1^ (Boyde & Firth, 2005; Firth et al., 2005) should take 60–90 days to infill from cement line (c. 100 μm radius) to finished canal (c. 10 μm radius) (Felder et al., 2017). Completion of remodelling at the organ scale relies on a number of secondary osteons working asynchronously in different locations and is likely to be complicated by individual variations in bone loading, training and resting patterns. Gentle preconditioning exercise that stimulates infilling and that does not suppress targeted remodelling is desirable, but the specific training regimens required to achieve this remain obscure.

Healed fractures are very common among mammals having appeared early in vertebrate evolution (Herbst et al., 2019), suggesting a survival advantage to risking fracture events because fractured bones have the capacity to heal well. We speculate that it may be a lesser cost to the organism during chronic high load periods to accumulate fatigue and defer remodelling with the concomitant fatigue fracture risk, than to incur the high metabolic cost of tissue turnover and repair.

While the most plausible biological explanation for the increased number and size of canals is associated with the remodelling process that occurs with low levels of mechanical loading, the association between duration of rest period and age is unknown. We selected a rest period of 4–16 weeks based on results of previous studies (Burr et al., 1989a, 1989b; McCarthy & Jeffcott, 1988) in which bone resorption was detected between 4 and 16 weeks.

To the authors’ knowledge, no previous studies have reported on the differences in fracture-free bone resorption between exercised and rested racehorses. Choosing non-fractured horses was paramount as horses with diagnosed catastrophic or non-catastrophic fractures may have had altered normal bone loading patterns and would not be truly representative of bone loading and resorption, although it was possible that other non-fracture related reasons may also have caused significant lameness that may have reduced the intensity of bone loading during high intensity exercise. Furthermore, this selection criterion resulted in a small sample size of 12 horses, which limits the power of the study to detect small differences between groups and increased the possibility of random error. Matching for age was not considered practical and would have further reduced case numbers; failure to do so here resulted in the exercised horses being on average younger than the rested horses (Table 1) (Holmes et al., 2014). We considered that restricting the selection criterion to high intensity exercise was appropriate because it has the advantage of reducing the confounding influence of training variation between horses, and is consistent with the design of other studies in the field (Holmes et al., 2014; Whitton et al., 2010, 2013). The loss of a tibia specimen was unfortunate because the two to three-fold difference in N.Ca/Ct.Ar and Tt.Ca.Ar/Ct.Ar for cutting cone-sized canals in the tibial site cannot be considered a robust result and is merely suggestive.

It was not possible to cut 100 μm sections from the entire cross section of the distal metaphysis of the third metacarpal bone and so our analysis was restricted to the lateral half of the bone in this location. The palmar/palmarolateral cortex of the third metacarpal bone is the most common location for stress fractures in the distal metaphyseal region of this bone (Shan et al., 2022) and, therefore, we believe that the results from this location are likely to reflect relevant findings. The anatomical location from which specimens from the tibia were sampled was chosen because this is a common location for stress fractures to occur in our population of horses based on diagnosis by nuclear scintigraphy. While others have reported the same finding in other jurisdictions (O’Sullivan & Lumsden, 2003), it is noteworthy that complete fracture of the tibia due to stress fracture is much more common in the proximal tibia (Samol et al., 2021). Consequently, intra-cortical resorption may be much more intense in bone of the proximal tibia than we determined from this study.

The numerical cutoffs that we used to classify canals into large and small groups are a simplification that aids in interpretation. The microradiographic data presented here do not capture the physiological status of each canal, only their sizes. We are also aware that a hard cutoff does not necessarily represent the diversity of modes by which resorption and infilling may occur and their dynamic nature, including substantial resting periods where neither resorption nor deposition are occurring. We believe that it is helpful to separate the very largest canals (Ca.Ar > 0.04 mm^2^; resorption space, cutting cone) from the moderately-sized (Ca.Ar > 0.002 mm^2^; infilling canal, closing cone) and smallest canals (Ca.Ar < 0.002 mm^2^; mature, Haversian canal) because these categories give some insight into the overall status of the bone and in particular whether porosity that can have resulted only from resorption is more or less common depending on the exercise or rested status of the animal.

Variation in bone geometry was minimised by cutting bone blocks in pre-determined standardised locations and visualising canals relative to the bone section’s centre of mass (Figure 5). The relationship between bone remodelling and bone geometry was not within the scope of this study but is likely to be affected by differences in mechanical loading (Davies, 2006; Nunamaker et al., 1989; Piotrowski et al., 1983), and changes to the second moment of area (Merritt & Davies, 2010; Nunamaker et al., 1990) to minimise peak bone strains. It is therefore not unreasonable to assume that different training regimens may have influenced the geometry of some of the bone sections, and the relationship between bone resorption and bone geometry should be explored in future studies. The counting of canals was automated with Fiji software which should have reduced bias. A thresholding technique was used for porosity segmentation but did contend with substantial variation in background intensity due to uneven sample thickness. Given the aim of this study, two-dimensional microradiography was considered a suitable imaging technique to identify bone resorption. However, future studies should use three dimensional imaging techniques such as X-ray microtomography and incorporate other markers of bone remodelling such as intra-vital calcein labelling to evaluate bone formation rate (Boyde & Firth, 2005).

## Conclusion

We found in limb bones, but not in ribs, that total canal area and the number of large canals, representing resorption activity, were greater in rested than in exercised animals, and that large canals were concentrated in the dorsal metacarpus, which is known to experience high strains during peak athletic loading. We surmise that the resorption phase of bone remodelling is significantly suppressed by high intensity exercise and is subsequently reactivated during a rest period to repair accumulated damage in a site-specific manner. During high intensity exercise periods, the balance between the mechanostat (increasing bone mass) and targeted remodelling (increased secondary osteonal cutting cone formation) appears to tilt towards the mechanostat. Targeted remodelling and the mechanostat may work synergistically in rest periods to remove damaged and excess bone tissue. Numerous large canals might exacerbate the risk of catastrophic failure should intense exercise be reintroduced prior to the osteoblastic infilling that completes the bone remodelling cycle. Future studies should be directed at determining appropriate training regimens to protect athletes’ bones when they return to exercise following an extended period of rest.

## Acknowledgements

We are grateful to have received funding from a City University of Hong Kong Postgraduate Studentship to RGCS (UGC-related research project 9610446, to MD), and from Arthritis Research UK to AB for the purchase of the Faxitron microradiography system.

## Conflict of interest

The authors have no conflict of interest.

## Authors’ contributions

Raniere Gaia Costa da Silva: formal analysis, methodology, writing original draft. Tsim Christopher Sun: methodology, writing original draft. Ambika Prasad Mishra: writing. Alan Boyde: obtained the grant for the Faxitron and performed the imaging. Michael Doube: supervision, writing original draft. Christopher Riggs: supervision, conceptualization, methodology, writing original draft. All authors contributed to review and editing and gave final approval for publication and agreed to be held accountable for the work performed therein.

## Supplementary material

TIFF images (8-bit) as used for analysis are available at doi:10.6084/m9.figshare.c.5910674. Full bit-depth (16-bit) DICOMs are available upon reasonable request. Bone statistics are available at doi:10.6084/m9.figshare.19410431. Pore information is available at doi:10.6084/m9.figshare.19410452. Full details of the image processing and statistical analysis are available at doi:10.6084/m9.figshare.19418933.

## Notes

https://doi.org/10.6084/m9.figshare.c.5910674

https://doi.org/10.6084/m9.figshare.19410431

https://doi.org/10.6084/m9.figshare.19410452

https://doi.org/10.6084/m9.figshare.19418933

### Competing Interest Statement

The authors have declared no competing interest.

### Summary of Updates

References to the humerus have been removed. Formatted for Journal of Anatomy.

## References

Allen, M.R., Iwata, K., Phipps, R. & Burr, D.B. (2006) Alterations in canine vertebral bone turnover, microdamage accumulation, and biomechanical properties following 1-year treatment with clinical treatment doses of risedronate or alendronate. Bone, 39, 872–879. Available from: https://doi.org/10.1016/j.bone.2006.04.028

Andreasen, C.M., Bakalova, L.P., Brüel, A., Hauge, E.M., Kiil, B.J., Delaisse, J.-M., et al. (2020) The generation of enlarged eroded pores upon existing intracortical canals is a major contributor to endocortical trabecularization. Bone, 130, 115127. Available from: https://doi.org/10.1016/j.bone.2019.115127

Andreasen, C.M., Delaisse, J.-M., van der Eerden, B.C., van Leeuwen, J.P., Ding, M. & Andersen, T.L. (2018) Understanding Age-Induced Cortical Porosity in Women: The Accumulation and Coalescence of Eroded Cavities Upon Existing Intracortical Canals Is the Main Contributor. Journal of Bone and Mineral Research, 33, 606–620. Available from: https://doi.org/10.1002/jbmr.3354

Armstrong, D.W., Rue, J.-P.H., Wilckens, J.H. & Frassica, F.J. (2004) Stress fracture injury in young military men and women. Bone, 35, 806–816. Available from: https://doi.org/10.1016/j.bone.2004.05.014

Bakalova, L.P., Andreasen, C.M., Thomsen, J.S., Brüel, A., Hauge, E.-M., Kiil, B.J., et al. (2018) Intracortical Bone Mechanics Are Related to Pore Morphology and Remodeling in Human Bone. Journal of Bone and Mineral Research, 33, 2177–2185. Available from: https://doi.org/10.1002/jbmr.3561

Behrens, J.C., Walker, P.S. & Shoji, H. (1974) Variations in strength and structure of cancellous bone at the knee. Journal of Biomechanics, 7, 201–207. Available from: https://doi.org/10.1016/0021-9290(74)90010-4

Bell, K.L., Loveridge, N., Power, J., Garrahan, N., Meggitt, B.F. & Reeve, J. (1999) Regional differences in cortical porosity in the fractured femoral neck. Bone, 24, 57–64. Available from: https://doi.org/10.1016/S8756-3282(98)00143-4

Boyde, A. (2003) The real response of bone to exercise. Journal of Anatomy, 203, 173–189. Available from: https://doi.org/10.1046/j.1469-7580.2003.00213.x

Boyde, A. & Firth, E.C. (2005) Musculoskeletal responses of 2-year-old Thoroughbred horses to early training. 8. Quantitative back-scattered electron scanning electron microscopy and confocal fluorescence microscopy of the epiphysis of the third metacarpal bone. New Zealand Veterinary Journal, 53, 123–32. Available from: https://doi.org/10.1080/00480169.2005.36489

Burr, D.B., Schaffler, M.B., Yang, K.H., Lukoschek, M., Sivaneri, N., Blaha, J.D., et al. (1989a) Skeletal change in response to altered strain environments: is woven bone a response to elevated strain? Bone, 10, 223–233. Available from: https://doi.org/10.1016/8756-3282(89)90057-4

Burr, D.B., Schaffler, M.B., Yang, K.H., Wu, D.D., Lukoschek, M., Kandzari, D., et al. (1989b) The effects of altered strain environments on bone tissue kinetics. Bone, 10, 215–221. Available from: https://doi.org/10.1016/8756-3282(89)90056-2

Busse, B., Hahn, M., Schinke, T., Püschel, K., Duda, G.N. & Amling, M. (2010) Reorganization of the femoral cortex due to age-, sex-, and endoprosthetic-related effects emphasized by osteonal dimensions and remodeling. Journal of Biomedical Materials Research Part A, 92A, 1440–1451. Available from: https://doi.org/10.1002/jbm.a.32432

Cappariello, A., Maurizi, A., Veeriah, V. & Teti, A. (2014) The Great Beauty of the osteoclast. Archives of Biochemistry and Biophysics, 558, 70–78. Available from: https://doi.org/10.1016/j.abb.2014.06.017

Carrier, T.K., Estberg, L., Stover, S.M., Gardner, I.A., Johnson, B.J., Read, D.H., et al. (1998) Association between long periods without high-speed workouts and risk of complete humeral or pelvic fracture in thoroughbred racehorses: 54 cases (1991-1994). Journal of the American Veterinary Medical Association, 212, 1582–1587.

Chalupa, R.L., Aberle, C. & Johnson, A.E. (2016) Observed Rates of Lower Extremity Stress Fractures After Implementation of the Army Physical Readiness Training Program at JBSA Fort Sam Houston. U.S. Army Medical Department Journal, 6–9.

Collins, C.J., Kozyrev, M., Frank, M., Andriotis, O.G., Byrne, R.A., Kiener, H.P., et al. (2019) The impact of age, mineralization, and collagen orientation on the mechanics of individual osteons from human femurs. Materialia, 100573. Available from: https://doi.org/10.1016/j.mtla.2019.100573

Davies, H.M.S. (2005) The timing and distribution of strains around the surface of the midshaft of the third metacarpal bone during treadmill exercise in one Thoroughbred racehorse. Australian Veterinary Journal, 83, 157–162. Available from: https://doi.org/10.1111/j.1751-0813.2005.tb11628.x

Davies, H.M.S. (2006) Estimating peak strains associated with fast exercise in Thoroughbred racehorses. Equine Veterinary Journal, 38, 383–386. Available from: https://doi.org/10.1111/j.2042-3306.2006.tb05573.x

Dempster, D.W., Compston, J.E., Drezner, M.K., Glorieux, F.H., Kanis, J.A., Malluche, H., et al. (2013) Standardized Nomenclature, Symbols, and Units for Bone Histomorphometry: A 2012 Update of the Report of the ASBMR Histomorphometry Nomenclature Committee. Journal of Bone and Mineral Research, 28, 2–17. Available from: https://doi.org/10.1002/jbmr.1805

Dittmer, K.E. & Firth, E.C. (2017) Mechanisms of bone response to injury. Journal of Veterinary Diagnostic Investigation, 29, 385–395. Available from: https://doi.org/10.1177/1040638716679861

Doube, M. (2022) Closing cones create conical lamellae in secondary osteonal bone. Royal Society Open Science, 9, 220712. Available from: https://doi.org/10.1098/rsos.220712

Felder, A.A., Phillips, C., Cornish, H., Cooke, M., Hutchinson, J.R. & Doube, M. (2017) Secondary osteons scale allometrically in mammalian humerus and femur. Royal Society Open Science, 4, 170431. Available from: https://doi.org/10.1098/rsos.170431

Firth, E.C., Rogers, C.W., Doube, M. & Jopson, N.B. (2005) Musculoskeletal responses of 2-year-old Thoroughbred horses to early training. 6. Bone parameters in the third metacarpal and third metatarsal bones. New Zealand Veterinary Journal, 53, 101–112. Available from: https://doi.org/10.1080/00480169.2005.36487

Frank, J.D., Ryan, M., Kalscheur, V.L., Ruaux-Mason, C.P., Hozak, R.R. & Muir, P. (2002) Aging and accumulation of microdamage in canine bone. Bone, 30, 201–206. Available from: https://doi.org/10.1016/S8756-3282(01)00623-8

Frost, H.M. (1969) Tetracycline-based histological analysis of bone remodeling. Calcified Tissue Research, 3, 211–237. Available from: https://doi.org/10.1007/bf02058664

Frost, H.M. (2003) Bone’s mechanostat: A 2003 update. Anatomical Record, 275A, 1081–1101. Available from: https://doi.org/10.1002/ar.a.10119

Gibson, V.A., Stover, S.M., Gibeling, J.C., Hazelwood, S.J. & Martin, R.B. (2006) Osteonal effects on elastic modulus and fatigue life in equine bone. Journal of Biomechanics, 39, 217–225. Available from: https://doi.org/10.1016/j.jbiomech.2004.12.002

Goldman, H.M., Bromage, T.G., Thomas, C.D.L. & Clement, J.G. (2003) Preferred collagen fiber orientation in the human mid-shaft femur. The Anatomical Record, 272A, 434–445. Available from: https://doi.org/10.1002/ar.a.10055

Goldman, H.M., Hampson, N.A., Guth, J.J., Lin, D. & Jepsen, K.J. (2014) Intracortical Remodeling Parameters Are Associated With Measures of Bone Robustness. The Anatomical Record, 297, 1817–1828. Available from: https://doi.org/10.1002/ar.22962

Herbst, E.C., Doube, M., Smithson, T.R., Clack, J.A. & Hutchinson, J.R. (2019) Bony lesions in early tetrapods and the evolution of mineralized tissue repair. Paleobiology, 45, 676–697. Available from: https://doi.org/10.1017/pab.2019.31

Hernandez, C.J. & Keaveny, T.M. (2006) A biomechanical perspective on bone quality. Bone, 39, 1173–1181. Available from: https://doi.org/10.1016/j.bone.2006.06.001

Holmes, J.M., Mirams, M., Mackie, E.J. & Whitton, R.C. (2014) Thoroughbred horses in race training have lower levels of subchondral bone remodelling in highly loaded regions of the distal metacarpus compared to horses resting from training. The Veterinary Journal, 202, 443–447. Available from: https://doi.org/10.1016/j.tvjl.2014.09.010

Hughes, J.M., Castellani, C.M., Popp, K.L., Guerriere, K.I., Matheny, R.W., Nindl, B.C., et al. (2020) The Central Role of Osteocytes in the Four Adaptive Pathways of Bone’s Mechanostat. Exercise and Sport Sciences Reviews, 48, 140–148. Available from: https://doi.org/10.1249/JES.0000000000000225

Hughes, J.M., Smith, M.A., Henning, P.C., Scofield, D.E., Spiering, B.A., Staab, J.S., et al. (2014) Bone formation is suppressed with multi-stressor military training. European Journal of Applied Physiology, 114, 2251–2259. Available from: https://doi.org/10.1007/s00421-014-2950-6

Ireland, A., Maden-Wilkinson, T., Ganse, B., Degens, H. & Rittweger, J. (2014) Effects of age and starting age upon side asymmetry in the arms of veteran tennis players: a cross-sectional study. Osteoporosis International, 25, 1389–1400. Available from: https://doi.org/10.1007/s00198-014-2617-5

Jacobs, J.M., Cameron, K.L. & Bojescul, J.A. (2014) Lower extremity stress fractures in the military. Clinics in Sports Medicine, 33, 591–613. Available from: https://doi.org/10.1016/j.csm.2014.06.002

Jaworski, Z.F. & Lok, E. (1972) The rate of osteoclastic bone erosion in Haversian remodeling sites of adult dog’s rib. Calcified Tissue Research, 10, 103–112. Available from: https://doi.org/10.1007/BF02012540

Komori, T. (2013) Functions of the osteocyte network in the regulation of bone mass. Cell and Tissue Research, 352, 191–198. Available from: https://doi.org/10.1007/s00441-012-1546-x

Lassen, N.E., Andersen, T.L., Pløen, G.G., Søe, K., Hauge, E.M., Harving, S., et al. (2017) Coupling of Bone Resorption and Formation in Real Time: New Knowledge Gained From Human Haversian BMUs. Journal of Bone and Mineral Research, 32, 1395–1405. Available from: https://doi.org/10.1002/jbmr.3091

Li, J., Mashiba, T. & Burr, D.B. (2001) Bisphosphonate Treatment Suppresses Not Only Stochastic Remodeling but Also the Targeted Repair of Microdamage. Calcified Tissue International, 69, 281–286. Available from: https://doi.org/10.1007/s002230010036

Martin, R.B. (2007) Targeted bone remodeling involves BMU steering as well as activation. Bone, 40, 1574–1580. Available from: https://doi.org/10.1016/j.bone.2007.02.023

Martin, R.B., Stover, S.M., Gibson, V.A., Gibeling, J.C. & Griffin, L.V. (1996) In vitro fatigue behavior of the equine third metacarpus: remodeling and microcrack damage analysis. Journal of Orthopaedic Research: Official Publication of the Orthopaedic Research Society, 14, 794–801. Available from: https://doi.org/10.1002/jor.1100140517

Mashiba, T., Hirano, T., Turner, C.H., Forwood, M.R., Johnston, C.C. & Burr, D.B. (2000) Suppressed bone turnover by bisphosphonates increases microdamage accumulation and reduces some biomechanical properties in dog rib. Journal of Bone and Mineral Research, 15, 613–620. Available from: https://doi.org/10.1359/jbmr.2000.15.4.613

McCalden, R.W., McGeough, J.A., Barker, M.B. & Court-Brown, C.M. (1993) Age-related changes in the tensile properties of cortical bone. The relative importance of changes in porosity, mineralization, and microstructure. The Journal of Bone and Joint Surgery. American Volume, 75, 1193–1205. Available from: https://doi.org/10.2106/00004623-199308000-00009

McCalden, R.W., McGeough, J.A. & Court-Brown, C.M. (1997) Age-related changes in the compressive strength of cancellous bone. The relative importance of changes in density and trabecular architecture. The Journal of Bone and Joint Surgery. American Volume, 79, 421–427. Available from: https://doi.org/10.2106/00004623-199703000-00016

McCarthy, R.N. & Jeffcott, L.B. (1988) Monitoring the effects of treadmill exercise on bone by non-invasive means during a progressive fitness programme. Equine Veterinary Journal. Supplement, 20, 88–92. Available from: https://doi.org/10.1111/j.2042-3306.1988.tb04653.x

McFarlin, S.C., Terranova, Carl.J., Zihlman, A.L., Enlow, D.H. & Bromage, T.G. (2008) Regional variability in secondary remodeling within long bone cortices of catarrhine primates: the influence of bone growth history. Journal of Anatomy, 213, 308–324. Available from: https://doi.org/10.1111/j.1469-7580.2008.00947.x

Merritt, J.S. & Davies, H.M.S. (2010) Metacarpal geometry changes during Thoroughbred race training are compatible with sagittal-plane cantilever bending: Cantilever bending of the equine metacarpus. Equine Veterinary Journal, 42, 407–411. Available from: https://doi.org/10.1111/j.2042-3306.2010.00209.x

Milgrom, C. & Finestone, A.S. (2017) The effect of stress fracture interventions in a single elite infantry training unit (1983-2015). Bone, 103, 125–130. Available from: https://doi.org/10.1016/j.bone.2017.06.026

Mori, S. & Burr, D.B. (1993) Increased intracortical remodeling following fatigue damage. Bone, 14, 103–109. Available from: https://doi.org/10.1016/8756-3282(93)90235-3

Nalla, R.K., Stölken, J.S., Kinney, J.H. & Ritchie, R.O. (2005) Fracture in human cortical bone: local fracture criteria and toughening mechanisms. Journal of Biomechanics, 38, 1517–1525. Available from: https://doi.org/10.1016/j.jbiomech.2004.07.010

Norman, T.L., Yeni, Y.N., Brown, C.U. & Wang, Z. (1998) Influence of microdamage on fracture toughness of the human femur and tibia. Bone, 23, 303–306. Available from: https://doi.org/10.1016/s8756-3282(98)00103-3

Nunamaker, D.M., Butterweck, D.M. & Provost, M.T. (1989) Some geometric properties of the third metacarpal bone: A comparison between the thoroughbred and standardbred racehorse. Journal of Biomechanics, 22, 129–134. Available from: https://doi.org/10.1016/0021-9290(89)90035-3

Nunamaker, D.M., Butterweck, D.M. & Provost, M.T. (1990) Fatigue fractures in thoroughbred racehorses: relationships with age, peak bone strain, and training. Journal of Orthopaedic Research, 8, 604–611.

O’Sullivan, C.B. & Lumsden, J.M. (2003) Stress fractures of the tibia and humerus in Thoroughbred racehorses: 99 cases (1992-2000). Journal of the American Veterinary Medical Association, 222, 491–8. Available from: https://doi.org/12597423

Parfitt, A.M. (2002) Targeted and nontargeted bone remodeling: relationship to basic multicellular unit origination and progression. Bone, 30, 5–7. Available from: https://doi.org/10.1016/S8756-3282(01)00642-1

Piotrowski, G., Sullivan, M. & Colahan, P.T. (1983) Geometric properties of equine metacarpi. Journal of Biomechanics, 16, 129–139. Available from: https://doi.org/10.1016/0021-9290(83)90036-2

Plotkin, L.I. (2014) Apoptotic osteocytes and the control of targeted bone resorption. Current Osteoporosis Reports, 12, 121–126. Available from: https://doi.org/10.1007/s11914-014-0194-3

Riggs, C.M., Lanyon, L.E. & Boyde, A. (1993a) Functional associations between collagen fibre orientation and locomotor strain direction in cortical bone of the equine radius. Anatomy and Embryology, 187, 231–8. Available from: https://doi.org/10.1007/BF00195760

Riggs, C.M. & Pilsworth, R. (2014) Repetitive strain injuries of the skeleton in high performance equine athletes. In: Hinchcliff, K.W., Kaneps, A.J., & Geor, R.J. (Eds.) Equine Sports Medicine and Surgery. Second Edition. Edinburgh: W.B. Saunders, pp. 457–471. Available online: https://doi.org/10.1016/B978-0-7020-4771-8.00022-3. Accessed 28 April 2022

Riggs, C.M., Vaughan, L.C., Evans, G.P., Lanyon, L.E. & Boyde, A. (1993b) Mechanical implications of collagen fibre orientation in cortical bone of the equine radius. Anatomy and Embryology, 187, 239–248. Available from: https://doi.org/10.1007/BF00195761

Riggs, C.M., Whitehouse, G.H. & Boyde, A. (1999) Pathology of the distal condyles of the third metacarpal and third metatarsal bones of the horse. Equine Veterinary Journal, 31, 140–148. Available from: https://doi.org/10.1111/j.2042-3306.1999.tb03807.x

Rueden, C.T., Schindelin, J., Hiner, M.C., DeZonia, B.E., Walter, A.E., Arena, E.T., et al. (2017) ImageJ2: ImageJ for the next generation of scientific image data. BMC Bioinformatics, 18, 529. Available from: https://doi.org/10.1186/s12859-017-1934-z

Samol, M.A., Uzal, F.A., Hill, A.E., Arthur, R.M. & Stover, S.M. (2021) Characteristics of complete tibial fractures in California racehorses. Equine Veterinary Journal, 53, 911–922. Available from: https://doi.org/10.1111/evj.13375

Schaffler, M.B. & Burr, D.B. (1988) Stiffness of compact bone: Effects of porosity and density. Journal of Biomechanics, 21, 13–16. Available from: https://doi.org/10.1016/0021-9290(88)90186-8

Schindelin, J., Arganda-Carreras, I., Frise, E., Kaynig, V., Longair, M., Pietzsch, T., et al. (2012) Fiji: an open-source platform for biological-image analysis. Nature Methods, 9, 676–682. Available from: https://doi.org/10.1038/nmeth.2019

Schneider, C.A., Rasband, W.S. & Eliceiri, K.W. (2012) NIH Image to ImageJ: 25 years of image analysis. Nature Methods, 9, 671–675. Available from: https://doi.org/10.1038/nmeth.2089

SciPy 1.0 Contributors, Virtanen, P., Gommers, R., Oliphant, T.E., Haberland, M., Reddy, T., et al. (2020) SciPy 1.0: fundamental algorithms for scientific computing in Python. Nature Methods, 17, 261–272. Available from: https://doi.org/10.1038/s41592-019-0686-2

Seref-Ferlengez, Z., Basta-Pljakic, J., Kennedy, O.D., Philemon, C.J. & Schaffler, M.B. (2014) Structural and mechanical repair of diffuse damage in cortical bone in vivo. Journal of Bone and Mineral Research, 29, 2537–2544. Available from: https://doi.org/10.1002/jbmr.2309

Seref-Ferlengez, Z., Kennedy, O.D. & Schaffler, M.B. (2015) Bone microdamage, remodeling and bone fragility: how much damage is too much damage? BoneKEy Reports, 4, 644. Available from: https://doi.org/10.1038/bonekey.2015.11

Shahar, R., Lukas, C., Papo, S., Dunlop, J.W.C. & Weinkamer, R. (2011) Characterization of the Spatial Arrangement of Secondary Osteons in the Diaphysis of Equine and Canine Long Bones. The Anatomical Record: Advances in Integrative Anatomy and Evolutionary Biology, 294, 1093–1102. Available from: https://doi.org/10.1002/ar.21405

Shan, R., Johnston, A.S., Rosanowski, S.M., O’Shea, J. & Riggs, C.M. (2022) Stress fracture of the palmar, distal cortex of the third metacarpal bone: A diagnostic challenge with a good prognosis. Equine Veterinary Journal, 54, 74–81. Available from: https://doi.org/10.1111/evj.13426

Shaw, C.N. & Ryan, T.M. (2012) Does skeletal anatomy reflect adaptation to locomotor patterns? cortical and trabecular architecture in human and nonhuman anthropoids. American Journal of Physical Anthropology, 147, 187–200. Available from: https://doi.org/10.1002/ajpa.21635

Shaw, C.N. & Stock, J.T. (2009) Intensity, repetitiveness, and directionality of habitual adolescent mobility patterns influence the tibial diaphysis morphology of athletes. American Journal of Physical Anthropology, 140, 149–159. Available from: https://doi.org/10.1002/ajpa.21064

Stover, S.M., Pool, R.R., Martin, R.B. & Morgan, J.P. (1992) Histological features of the dorsal cortex of the third metacarpal bone mid-diaphysis during postnatal growth in thoroughbred horses. Journal of Anatomy, 181 (Pt 3), 455–469.

The Hong Kong Jockey Club (2021) Racing Information Database. The Hong Kong Jockey Club. Available online: https://racing.hkjc.com/racing/information/english/Horse/SelectHorse.aspx

Turnbull, T.L., Baumann, A.P. & Roeder, R.K. (2014) Fatigue microcracks that initiate fracture are located near elevated intracortical porosity but not elevated mineralization. Journal of Biomechanics, 47, 3135–3142. Available from: https://doi.org/10.1016/j.jbiomech.2014.06.022

Ural, A. & Vashishth, D. (2007) Effects of intracortical porosity on fracture toughness in aging human bone: a microCT-based cohesive finite element study. Journal of Biomechanical Engineering, 129, 625–631. Available from: https://doi.org/10.1115/1.2768377

Vallance, S.A., Entwistle, R.C., Hitchens, P.L., Gardner, I.A. & Stover, S.M. (2013) Case-control study of high-speed exercise history of Thoroughbred and Quarter Horse racehorses that died related to a complete scapular fracture: Exercise history of racehorses with a catastrophic scapular fracture. Equine Veterinary Journal, 45, 284–292. Available from: https://doi.org/10.1111/j.2042-3306.2012.00644.x

Wachter, N.J., Krischak, G.D., Mentzel, M., Sarkar, M.R., Ebinger, T., Kinzl, L., et al. (2002) Correlation of bone mineral density with strength and microstructural parameters of cortical bone in vitro. Bone, 31, 90–95. Available from: https://doi.org/10.1016/S8756-3282(02)00779-2

Whitton, R.C., Mirams, M., Mackie, E.J., Anderson, G.A. & Seeman, E. (2013) Exercise-induced inhibition of remodelling is focally offset with fatigue fracture in racehorses. Osteoporosis International, 24, 2043–2048. Available from: https://doi.org/10.1007/s00198-013-2291-z

Whitton, R.C., Trope, G.D., Ghasem-Zadeh, A., Anderson, G.A., Parkin, T.D.H., Mackie, E.J., et al. (2010) Third metacarpal condylar fatigue fractures in equine athletes occur within previously modelled subchondral bone. Bone, 47, 826–831. Available from: https://doi.org/10.1016/j.bone.2010.07.019

Wik, E.H., Lolli, L., Chamari, K., Materne, O., Salvo, V.D., Gregson, W., et al. (2020) Injury patterns differ with age in male youth football: a four-season prospective study of 1111 time-loss injuries in an elite national academy. British Journal of Sports Medicine [Preprint]. Available from: https://doi.org/10.1136/bjsports-2020-103430

Yeni, Y.N., Brown, C.U., Wang, Z. & Norman, T.L. (1997) The influence of bone morphology on fracture toughness of the human femur and tibia. Bone, 21, 453–459. Available from: https://doi.org/10.1016/S8756-3282(97)00173-7

Zack, G.W., Rogers, W.E. & Latt, S.A. (1977) Automatic measurement of sister chromatid exchange frequency. Journal of Histochemistry & Cytochemistry, 25, 741–753. Available from: https://doi.org/10.1177/25.7.70454

